# Serum Electrolyte Concentrations Are Associated With Motor Nerve Conduction Velocity in Children

**DOI:** 10.64898/2026.10.07.757353

**Authors:** Philip J. Broser

## Abstract

Axonal action potential propagation critically depends on tightly regulated intra- and extracellular electrolyte concentrations and acid–base homeostasis. Experimental studies have demonstrated that changes in extracellular sodium, potassium, and calcium concentrations alter axonal excitability and conduction properties. However, it remains unknown whether small variations in electrolyte concentrations within the physiological range are associated with measurable changes in nerve conduction velocity in humans.

In this study, data from neurologically healthy children who underwent both nerve conduction studies and blood gas analysis were analyzed to investigate the relationship between electrolyte concentrations and median motor nerve conduction velocity. Nerve conduction velocity was significantly negatively correlated with ionized calcium concentration and positively correlated with sodium concentration, whereas no significant correlation was observed for potassium. In an a priori defined multiple linear regression model, ionized calcium, sodium, and bicarbonate remained significantly associated with nerve conduction velocity after correction for multiple testing.

These findings indicate that even small variations in electrolyte concentrations within the physiological range are associated with measurable differences in peripheral nerve conduction velocity and suggest an additional contribution of acid–base status to nerve conduction.

## Introduction

Axonal action potential propagation critically depends on tightly regulated intra- and extracellular electrolyte concentrations and acid–base homeostasis. In particular, the concentrations of sodium (Na⁺), potassium (K⁺), and calcium (Ca²⁺) within the extracellular, periaxonal, and nodal/paranodal compartments influence membrane potential, ion-channel function, and the efficient propagation of action potentials along myelinated axons (Poliak and Peles, 2003; Freeman et al., 2016; Rasband and Peles, 2016; Schwarz, 2021).

Previous studies in adults with renal and metabolic disorders have demonstrated associations between systemic electrolyte disturbances and peripheral nerve function. Alterations in calcium homeostasis have been associated with changes in nerve conduction velocity (NCV) and peripheral nerve function (Goldstein et al., 1978; Diniz et al., 2013). Similarly, associations between serum potassium and sodium concentrations and electrophysiological measures of peripheral nerve function have been reported (Ørskov et al., 1994; Kiernan et al., 2002). Importantly, some of these associations have been observed across relatively modest variations in electrolyte concentrations (Ørskov et al., 1994), raising the question of whether changes within or close to the physiological range may already exert measurable effects on peripheral nerve conduction.

The relationship between circulating electrolyte concentrations and the extracellular environment of neural tissue differs fundamentally between the central and peripheral nervous systems. In the central nervous system (CNS), the blood–brain and blood– cerebrospinal fluid barriers tightly regulate the ionic composition of the neuronal extracellular environment (Husted and Reed, 1976; Saunders et al., 2016). This regulation is particularly pronounced for potassium, for which cerebrospinal fluid concentrations remain remarkably stable despite variations in plasma potassium (Husted and Reed, 1976; Saunders et al., 2016). Sodium concentrations in the CSF are also subject to active regulation and can vary independently of plasma sodium under some physiological and pathological conditions (Harrington et al., 2006).

The situation differs in the peripheral nervous system (PNS). Peripheral axons are protected by the blood–nerve barrier, formed primarily by tight junctions between endoneurial endothelial cells and by the perineurium (Ubogu, 2013; Palladino et al., 2017). Although the blood–nerve barrier is highly restrictive to proteins and many hydrophilic solutes, experimental studies demonstrate measurable permeability of endoneurial capillaries to small inorganic ions including K⁺, Na⁺, and Cl⁻ (Weerasuriya, 1987). Direct measurements have demonstrated characteristic concentrations of sodium, potassium, and chloride within endoneurial fluid (Myers et al., 1983). Importantly, experimentally induced changes in circulating potassium can be transmitted through the endoneurial capillary wall and rapidly influence the extracellular environment of the paranodal region (Weerasuriya, 1987). Thus, although the endoneurial ionic environment is regulated, peripheral axons may be more directly exposed to changes in circulating small-ion concentrations than neurons within the CNS. Evidence for calcium is less extensive, and direct quantitative data linking circulating ionized calcium to endoneurial calcium concentrations remain limited.

Glucose is also transported across the blood–nerve barrier by facilitated transport mechanisms. Glucose transporters are expressed in both endoneurial microvascular endothelial cells and perineurial cells, supporting an important role of circulating glucose as an energy substrate for peripheral nerves (Rechthand et al., 1989; Gerhart and Drewes, 1990). Acute changes in glucose concentration do not necessarily produce corresponding immediate changes in NCV; in healthy adults, short-term hyperglycemia itself had little detectable effect on conduction velocity once concomitant metabolic and electrolyte changes were considered (Ørskov et al., 1994). In contrast, chronic glycemic exposure and glucose variability are associated with impaired peripheral nerve conduction in diabetes, including in children and adolescents (Oberhauser et al., 2023).

Acid–base homeostasis represents an additional potential determinant of axonal function. Carbon dioxide readily crosses biological membranes, providing a mechanism by which systemic changes in pCO₂ may influence tissue acid–base conditions. Experimental manipulation of extracellular pH in myelinated peripheral axons alters excitability through effects on several ion conductances, including persistent sodium currents and potassium currents (Maurer et al., 2012). Systemic acid–base status may therefore represent an additional source of physiological variability in peripheral nerve electrophysiology.

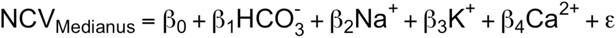

These observations raise the question of whether relatively small variations in systemic electrolyte concentrations and acid–base status are associated with measurable changes in peripheral nerve conduction, even in the absence of overt electrolyte or metabolic disturbances. Previous experimental and human data suggest that such short-term effects are biologically plausible (Ørskov et al., 1994; Weerasuriya, 1987; Kiernan et al., 2002; Maurer et al., 2012), but their magnitude and consistency within a predominantly healthy pediatric population have not been established.

Advances in nerve conduction studies allow differences in the conduction properties of large myelinated peripheral nerve fibers to be quantified with high precision.

Interpretation requires consideration of physiological variables, particularly age and maturation and precise control of the temperature during measurement (Parano et al., 1993; García et al., 2000). Investigating subtle physiological influences on nerve conduction in adults may additionally be complicated by cumulative exposure to metabolic, vascular, renal, and neurological disease.

The pediatric population provides a particularly valuable setting in which to investigate these relationships. Peripheral nerve conduction undergoes substantial maturation during infancy and childhood. NCV is considerably lower around birth, increases rapidly during the first years of life, and approaches adult values during early childhood (Parano et al., 1993; García et al., 2000; Farrar et al., 2013). Consequently, rigorous adjustment for age and developmental stage is essential in pediatric nerve conduction studies.

At our tertiary pediatric center, children with a broad spectrum of neurological disorders are evaluated in a specialized pediatric clinical neurophysiology and neuromuscular unit. A substantial subgroup consists of children undergoing assessment for traumatic peripheral nerve injuries. To avoid overlooking an underlying hereditary or generalized neuropathy in these patients, we established a standardized electrophysiological screening protocol that includes examination of the clinically unaffected contralateral side. The protocol comprises motor nerve conduction studies of the tibial, peroneal, and median nerves and sensory nerve conduction studies of the median nerve. The same combination of nerve conduction measurements has previously been used in our studies of pediatric peripheral nerve function (Oberhauser et al., 2023).

Many of these children undergo blood sampling as part of their clinical evaluation. This routinely includes blood-gas analysis with simultaneous measurement of acid–base parameters and electrolytes. This setting provides an opportunity to investigate the association between systemic electrolyte and acid–base homeostasis and peripheral nerve conduction in a pediatric population without known chronic systemic or neurological disease.

The aim of the present study was therefore to determine whether variations in serum sodium, potassium, and ionized calcium concentrations and systemic acid–base parameters are associated with nerve conduction velocity in children. In this study we specifically focus on the motor conduction velocity of the median nerve, which can be measured with highest pression across a large age range (Oberhauser et al., 2023).

## Methods

### Study population

A total of 41 children were screened for eligibility. All screened participants had been referred for evaluation of a unilateral traumatic peripheral nerve injury. Injuries were typically associated with fractures of the humerus or forearm following low-energy trauma, such as falls from low height or low-velocity bicycle accidents. Children with concomitant head injury, major polytrauma, or injuries affecting the contralateral extremities were not considered eligible.

Electrophysiological assessment was performed after the acute phase of the injury, typically 7–10 days after the traumatic event. This corresponds to the time at which a persisting peripheral nerve injury is commonly identified clinically and referred for specialized neurophysiological evaluation.

Of the 41 children screened, 33 (15 female) fulfilled the inclusion criteria and were included in the final analysis. Participants were excluded if the interval between nerve conduction studies and laboratory sampling exceeded 2 hours, informed consent was unavailable, relevant laboratory or electrophysiological measurements were missing, a chronic systemic or neurological disorder was present, or there was a family history suggestive of a hereditary neurological disorder.

The study was conducted as part of an ongoing research project approved by the responsible local ethics committee (2022-00216, EKOS 22/018). Written informed consent was obtained in accordance with the approved study protocol.

### Nerve conduction studies

All included children underwent a standardized peripheral nerve conduction protocol. Motor nerve conduction studies were performed for the peroneal, tibial, and median nerves, and sensory nerve conduction studies were performed for the median nerve. For the purpose of the present analysis, only the measurement of the median nerve on the unaffected side was included.

All examinations were performed according to the standardized St. Gallen pediatric neuropathy screening protocol, previously applied to studies of peripheral nerve function in children and adolescents (Oberhauser et al., 2023). Part of this protocol is to give the participant sufficient time to accommodate to the constant controlled temperature in the neurophysiology laboratory. Age-dependent interpretation is particularly important in children because peripheral nerve conduction undergoes marked maturation during the first years of life (Parano et al., 1993; García et al., 2000; Farrar et al., 2013). Therefore, only children above the age of two were included. For all neurophysiology measurements the Sierra Summit System (Cadwell Industries, Inc., Kennewick, WA, USA) was used.

### Laboratory measurements

Following the neurophysiological examination, participants underwent capillary blood sampling as part of their clinical assessment. The interval between electrophysiological examination and blood sampling was limited to a maximum of 2 hours. For blood gas analysis the cobas b 123 POC system (Roche Diagnostics International Ltd, Rotkreuz, Switzerland) was used.

Capillary blood-gas analysis included measurements of acid–base status and electrolytes, including pH, partial pressure of carbon dioxide (pCO₂), bicarbonate, base excess, sodium (Na⁺), potassium (K⁺), total calcium, and ionized calcium (Ca²⁺). Actual bicarbonate concentration (cHCO₃⁻) was calculated by the cobas POC system from measured pH and pCO₂ according to

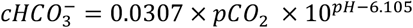

as recommended by the Clinical and Laboratory Standards Institute (CLSI C46-A2). All samples were obtained and analyzed using the hospital’s routine clinical laboratory infrastructure.

### Data acquisition and management

All clinical, neurophysiological, ultrasound, and laboratory measurements were acquired using the hospital’s routine clinical infrastructure. A dedicated study participant list was maintained on a secured research computer in accordance with institutional data-protection requirements.

For the present analysis, nerve conduction and laboratory data were extracted from the respective clinical information systems and merged into a single study dataset. Prior to statistical analysis, data were checked for completeness, plausibility, and consistency.

### Statistical analysis

All statistical analyses were performed in R (Version 4.4.1). Continuous variables were summarized using appropriate descriptive statistics according to their distributions. The relationships between sodium, potassium and calcium and median motor nerve conduction velocity were initially explored graphically using scatter plots. Pairwise correlations between electrophysiological and laboratory variables were subsequently calculated using Pearson’s correlation coefficients. To account for multiple testing, p-values were adjusted using the Holm method (Holm, 1979), which controls the family-wise error rate.

Based on these exploratory analyses and predefined physiological considerations, a linear regression models was constructed to investigate the association between electrolyte and acid–base parameters and nerve conduction velocity. The model assumptions were assessed by inspection of residual distributions and diagnostic plots and by the findings of the experimental study from (Frankenhaeuser, 1957).

The following model was a priori defined and tested in the study:

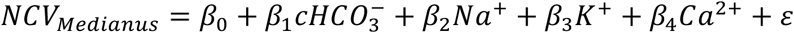

## Results

### Demographics and data

A total of 33 children (15 female) were included in the study. The demographic characteristics of the study population are summarized in Table 1. As specified in the Methods section, the minimum age for inclusion was 2 years. By this age, peripheral nerve myelination is already well advanced, and nerve conduction velocity has reached more than 80% of adult values. Table 2 summarizes the distribution of the measured variables and the number of observations available for each parameter.

**Table 1.** Participant characteristics.

|  | N | Mean | SD | Q1 | Median | Q3 | Min | Max |
| --- | --- | --- | --- | --- | --- | --- | --- | --- |
| Age | 33 | 10.7 | 4.3 | 6.9 | 10.6 | 14.1 | 2.2 | 17.4 |
| Height | 31 | 145.8 | 24.7 | 125.2 | 151.4 | 161.5 | 94.0 | 196.0 |
| Weight | 32 | 38.4 | 17.1 | 25.1 | 35.3 | 52.7 | 12.9 | 84.0 |

Values are presented as mean, standard deviation, median, interquartile range, and range.

**Table 2.** Data characteristics.

|  | N | Mean | SD | Q1 | Median | Q3 | Min | Max |
| --- | --- | --- | --- | --- | --- | --- | --- | --- |
| Median motor nerve conduction velocity | 33 | 56.69 | 4.57 | 53.00 | 56.00 | 60.00 | 47.00 | 67.00 |
| Median motor NCV Z-score | 33 | -0.10 | 1.05 | -0.91 | -0.32 | 0.74 | -1.61 | 2.32 |
| Skin temperature | 31 | 32.68 | 1.22 | 32.00 | 32.80 | 33.30 | 30.40 | 35.20 |
| Actual bicarbonate | 32 | 21.22 | 1.65 | 20.17 | 20.80 | 22.33 | 18.10 | 25.00 |
| Sodium | 33 | 139.95 | 1.79 | 139.00 | 140.00 | 141.00 | 135.00 | 143.00 |
| Potassium | 33 | 4.38 | 0.45 | 4.00 | 4.30 | 4.70 | 3.90 | 5.80 |
| Calcium | 33 | 1.19 | 0.04 | 1.17 | 1.18 | 1.20 | 1.12 | 1.26 |

NCV, nerve conduction velocity.

### Scatter plot

For an initial assessment of the relationships between electrolyte concentrations and nerve conduction velocity, the data were visualized using scatter plots (Figure 1). Median motor nerve conduction velocity was plotted against sodium (Panel A), potassium (Panel B), and ionized calcium concentrations (Panel C). Pearson correlation coefficients were calculated for all three electrolytes.

**Figure 1.**
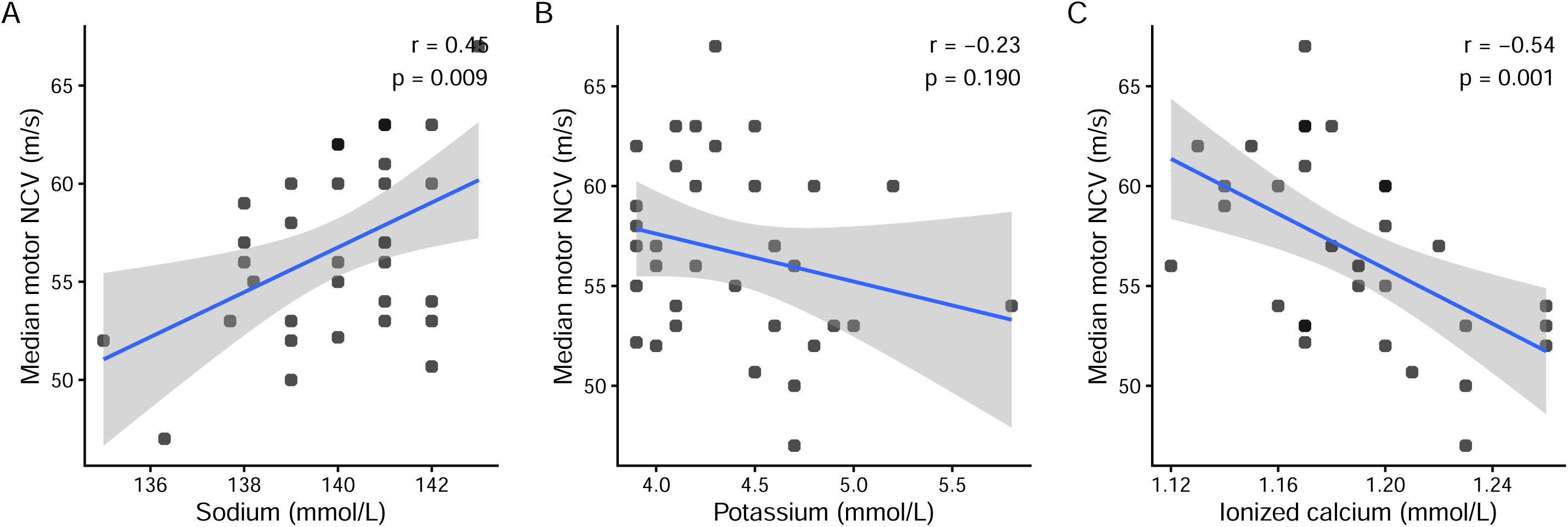
Association between electrolytes and median motor NCV.

Median motor nerve conduction velocity was significantly positively correlated with sodium concentration (r = 0.45, 95% CI [0.12 to 0.68], p = 0.009) and significantly negatively correlated with ionized calcium concentration (r = -0.54, 95% CI [-0.75 to - 0.24], p = 0.001), whereas no significant correlation was observed for potassium (r = - 0.23, 95% CI [-0.53 to 0.12], p = 0.190).

The direction of the associations with sodium and calcium is consistent with the electrophysiological effects of extracellular ion concentrations described by Frankenhaeuser (1957). Within the investigated concentration range, increasing sodium concentrations were associated with higher nerve conduction velocities, whereas increasing ionized calcium concentrations were associated with lower nerve conduction velocities. Potassium showed a tendency toward lower nerve conduction velocities at higher concentrations; however, this association was weak and did not reach statistical significance.

### Multiple linear regression modell

To account for the simultaneous effects of the different ions on nerve conduction velocity, a multiple linear regression model was defined a priori:

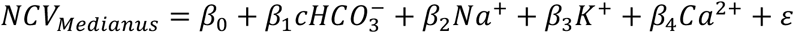

The prespecified model was fitted to the recorded data, and the results of the multiple linear regression analysis are presented in Table 3 and graphically summarized as a forest plot in Figure 2. After adjustment for multiple testing using the Holm method, the associations of bicarbonate, sodium, and ionized calcium with nerve conduction velocity remained statistically significant (Holm-adjusted p-values: bicarbonate, p = 0.0276; sodium, p = 0.0200; potassium, p = 0.4780; ionized calcium, p = 0.0276).

**Figure 2.**
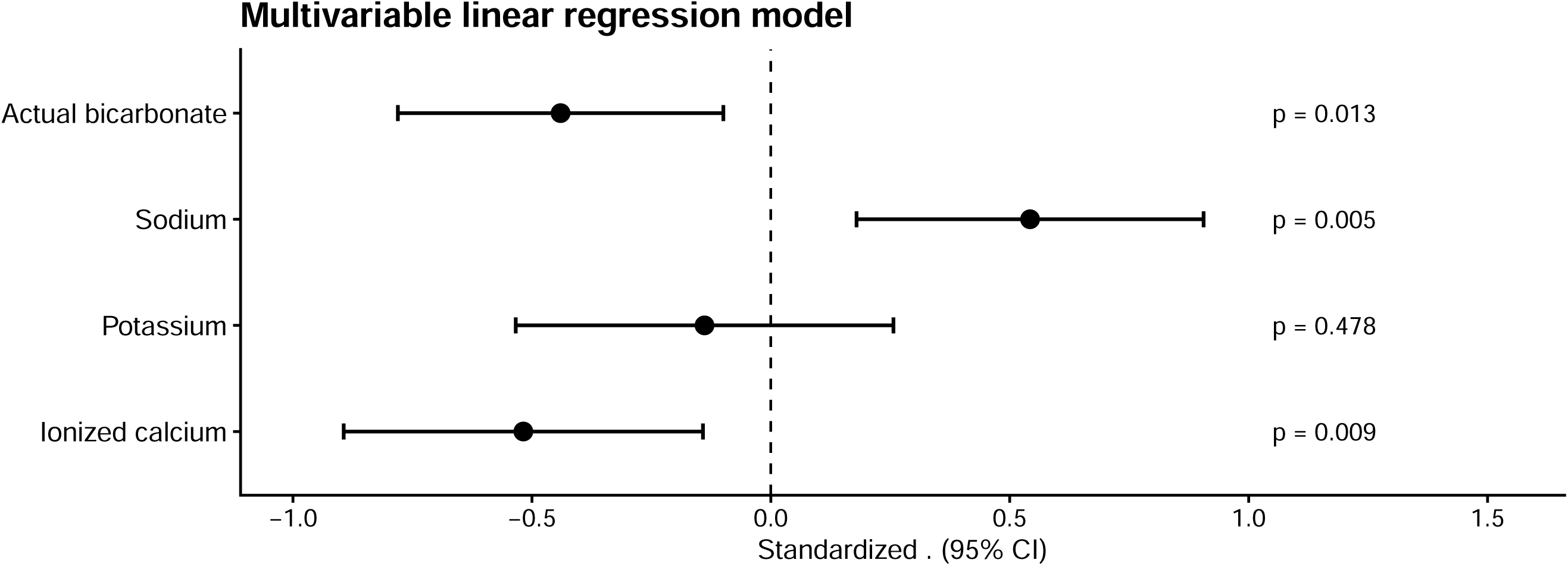
Multivariable linear regression model.

**Table 3.** Multivariable linear regression analysis.

| Predictor | Estimate | 95% CI | Standardized $\beta$ | SE | t | p |
| --- | --- | --- | --- | --- | --- | --- |
| Intercept | -32.00 | -168.95 to 104.95 | — | 66.75 | -0.48 | 0.636 |
| Actual bicarbonate | -1.14 | -2.02 to -0.26 | -0.44 | 0.43 | -2.65 | 0.013 |
| Sodium | 1.38 | 0.46 to 2.30 | 0.54 | 0.45 | 3.07 | 0.005 |
| Potassium | -1.32 | -5.09 to 2.44 | -0.14 | 1.84 | -0.72 | 0.478 |
| Ionized calcium | -62.36 | -107.64 to -17.09 | -0.52 | 22.07 | -2.83 | 0.009 |

N = 32; $R^2 = 0.486$ ; adjusted $R^2 = 0.410$ ; residual SE = 3.295; $F(4, 27) = 6.38$ ; model $p < 0.001$ . Standardized $\beta$ represents the change in outcome SD per 1-SD increase in the predictor.

As illustrated by the standardized regression coefficients in Figure 2, the individual ions showed associations of different directions with nerve conduction velocity. Higher ionized calcium concentrations were independently associated with lower nerve conduction velocities, whereas higher sodium concentrations were associated with higher nerve conduction velocities. Bicarbonate also showed a significant independent association with nerve conduction velocity, while no significant association was observed for potassium.

## Discussion

Previous human studies, particularly in patients with renal failure and other metabolic disorders, together with extensive experimental evidence, have demonstrated that physiological action potential propagation in peripheral nerves depends on tightly regulated extracellular and intracellular ion concentrations (Goldstein et al., 1978; Poliak and Peles, 2003; Freeman et al., 2016; Rasband and Peles, 2016). In chronic renal failure, abnormalities of axonal excitability correlate closely with serum potassium, and reduction of potassium during hemodialysis reverses electrophysiological features of axonal depolarization (Kiernan et al., 2002). While pronounced electrolyte disturbances are known to alter peripheral nerve excitability and conduction, considerably less is known about whether relatively small variations within or close to the physiological range are associated with measurable differences in nerve conduction. Importantly, however, the latter concept is not entirely unprecedented: Ørskov et al. (1994) demonstrated small changes in median NCV in healthy adults occurring in parallel with relatively modest changes in serum sodium and potassium concentrations.

In the present study, peripheral nerve conduction for the motor part of the median nerve was examined in children without known chronic systemic or neurological disease and related to electrolyte and acid–base parameters obtained from temporally closely matched blood samples. Electrophysiological measurements were performed using a standardized pediatric nerve conduction protocol that has previously been applied in studies of peripheral nerve function in children, including investigations of diabetic neuropathy (Oberhauser et al., 2023).

Despite the relatively small sample size, we identified consistent associations between several circulating electrolyte concentrations and NCV. The most prominent and consistent association was observed for ionized calcium and the plasma sodium concentrations.

At first sight, the prominent association with calcium may appear surprising, as rapid action potential propagation in myelinated axons is primarily dependent on voltage-gated sodium channels concentrated at the node of Ranvier, while potassium channels located within nodal and juxtaparanodal domains contribute to membrane potential and repolarization (Poliak and Peles, 2003; Freeman et al., 2016; Rasband and Peles, 2016; Schwarz, 2021). Calcium nevertheless has important modulatory roles in axonal physiology. Zhang and David (2016) demonstrated activity-dependent Ca²⁺ influx localized to nodal and perinodal regions of normally myelinated peripheral motor axons, involving T-type calcium channels and reverse-mode Na⁺/Ca²⁺ exchange. Experimental alteration of extracellular calcium also influences peripheral nerve excitability and conduction (Mert et al., 2003). Furthermore, experimental studies of uremia demonstrated that increased peripheral nerve calcium was associated with markedly reduced motor nerve conduction velocity (Goldstein et al., 1978). These observations provide biological plausibility for an association between calcium homeostasis and peripheral nerve conduction, although the present observational data cannot establish a causal relationship.

An associations was also observed for the potassium concentrations and NCV. Notably, the direction of this associations was as expected reverse to the effect of the sodium ions.

To summartize the observation sfrom this study that an increase in calcium lowers the NCV and an increase in sodium accelartes the NCV and the study favors a negative effect of an increase of the potassium concentration on the NCV.

This finding is compatible with the experimental observations that extracellular potassium strongly influences the axonal resting membrane potential, and human studies demonstrate axonal depolarization with increasing serum potassium (Kiernan et al., 2002). Sodium, in contrast, determines the electrochemical driving force for the inward current responsible for nodal depolarization (Poliak and Peles, 2003; Freeman et al., 2016; Rasband and Peles, 2016).

The net relationship between circulating electrolyte concentrations and measured NCV is therefore likely to reflect interactions among resting membrane potential, sodium-channel availability, nodal and internodal conductances, and the extracellular ionic environment. Given the exploratory nature and limited sample size of the present study, detailed mechanistic interpretation of the direction of these associations would be premature.

Furthermore, other biologically relevant ions were not available in the present dataset or were not studied due to the limited sizte of the dataset. In particular, phosphate and magnesium were not routinely measured. Both may influence peripheral nerve physiology. Experimental and clinical studies have demonstrated effects of extracellular magnesium on nerve conduction (Fleming et al., 1972; Mert et al., 2003), and associations between magnesium, phosphate, and electrophysiological measures of peripheral nerve function have also been reported in metabolic disease (Zhang et al., 2018). Future prospective studies should therefore include these parameters.

The observed associations are particularly relevant in the context of the blood–nerve barrier. The blood–nerve barrier is formed by tight-junction-bearing endoneurial endothelial cells together with the perineurial diffusion barrier and provides substantial restriction of solute exchange between blood and endoneurium (Ubogu, 2013; Palladino et al., 2017). It should therefore not be regarded as freely permeable. However, small inorganic ions represent an important exception to a simplistic barrier model.

Weerasuriya (1987) directly measured permeability of endoneurial capillaries to K⁺, Na⁺, and Cl⁻ and demonstrated substantial permeability to all three ions. Importantly, vascular perfusion with elevated potassium produced conduction block, and the kinetics indicated that changes in endoneurial K⁺ were transmitted to the paranodal extracellular space (Weerasuriya, 1987). Direct measurements also confirm the presence of characteristic Na⁺, K⁺, and Cl⁻ concentrations within endoneurial fluid (Myers et al., 1983). These observations provide a plausible physiological link between circulating electrolyte concentrations and the extracellular ionic environment encountered by peripheral axons. Direct evidence for equivalent serum-to-endoneurium coupling of calcium remains considerably more limited.

Acid–base status represents an additional potential modifier of peripheral nerve physiology. Experimental manipulation of extracellular pH in myelinated peripheral axons produces systematic changes in excitability. Increasing proton concentration increases threshold and reduces persistent sodium current, with additional effects on potassium conductances (Maurer et al., 2012). Human studies in critical illness have likewise demonstrated relationships between systemic metabolic changes and axonal membrane excitability (Z’Graggen et al., 2006).

In the present study, the actual bicarbonate concentration was used as a marker for the acid–base status. The actual bicarbonate is a calculated parameter from pH, pCO and should therefore interpreted cautiously.

Several limitations of this study need to be considered. Most importantly, the sample size was relatively small, particularly after selecting only children older than two years. The findings should therefore be regarded as exploratory and require confirmation in larger independent cohorts.

Especially the younger age group below the age of two should be studied in great detail. With Myelination the ion chanel distribution along the axon significantly alters.

Especially the distribution of potassium channels in the perinodal area shifts to the adult configuration and so it is expected that the effect of changes of the serum electrolytes have an distinct effect which should bei further analyzed.

Furthermore, although the participants had no known chronic systemic or neurological disorders, they cannot be considered completely healthy controls. All children had sustained a traumatic injury approximately one to two weeks before electrophysiological assessment. Acute and subacute physiological responses to trauma could potentially have influenced the measured systemic parameters. Whether such injury also modifies the endoneurial microenvironment of clinically unaffected contralateral nerves is unknown. This possibility cannot be excluded by the present study and should be considered when interpreting the findings.

An additional limitation is that circulating concentrations were used as surrogate measures of the extracellular environment of peripheral nerves. Although experimental evidence demonstrates passage of Na⁺, K⁺, and Cl⁻ across endoneurial capillaries (Weerasuriya, 1987) and direct measurements characterize the ionic composition of endoneurial fluid (Myers et al., 1983), serum concentrations cannot be assumed to equal endoneurial concentrations. This limitation is particularly relevant for calcium, for which direct quantitative evidence relating serum ionized calcium to endoneurial calcium remains limited.

Nevertheless, obtaining paired electrophysiological and laboratory measurements from children without chronic disease is inherently difficult. Prospective studies involving nerve conduction studies and blood sampling solely for research purposes raise practical and ethical challenges, particularly in younger children. The present study takes advantage of clinically indicated investigations to examine physiological relationships that would otherwise be difficult to study in a pediatric population.

The present findings also have potential methodological implications for clinical neurophysiology. Age and temperature are established determinants of peripheral nerve conduction and are important considerations in the interpretation of pediatric nerve conduction studies (Parano et al., 1993; García et al., 2000). Our results raise the possibility that electrolyte and acid–base status represent additional sources of physiological variability. Whether measurement of these parameters should become part of routine nerve conduction assessment cannot be determined from the present exploratory study. However, their documentation should be considered in future studies investigating subtle differences in peripheral nerve conduction, particularly when metabolic or systemic influences are of interest.

These observations may also be relevant for studies of metabolic neuropathies. In type 1 diabetes, chronic glycemic exposure and glucose variability are associated with reduced NCV, including in children and adolescents (Oberhauser et al., 2023). At the same time, acute experimental studies suggest that short-term hyperglycemia per se has little immediate effect on NCV and that concomitant electrolyte and hormonal changes may contribute to short-term alterations (Ørskov et al., 1994). A better understanding of the short-term physiological effects of electrolytes, acid–base status, and other circulating factors may therefore help distinguish acute metabolic influences on nerve conduction from structural or chronic metabolic changes of the peripheral nerve.

In conclusion, this exploratory study provides evidence that variations in circulating electrolyte concentrations within a relatively narrow clinical range are associated with measurable differences in peripheral nerve conduction in children. Calcium and Sodium showed the most consistent association, while associations were also observed for potassium, and the study shows that the acid-base status should be taken into consideration when analyzing nerve conduction studies.

Previous experimental and human data support the biological plausibility of effects of these ions on peripheral axonal excitability and conduction (Goldstein et al., 1978; Fleming et al., 1972; Ørskov et al., 1994; Weerasuriya, 1987; Kiernan et al., 2002; Mert et al., 2003; Maurer et al., 2012; Zhang and David, 2016). The present findings extend these observations to a pediatric population without known chronic systemic or neurological disease. Larger prospective studies are required to confirm these associations, quantify their effect sizes, determine whether they remain independently associated with NCV after rigorous adjustment for developmental and technical covariates, and clarify their underlying mechanisms and clinical relevance.

## Acknowledgements

I thank the members of the FND team for their contribution to the electrophysiological measurements. Special thank to Dr. Erin West for her critical review of the manuscript and review of the statistical analyses and a big thanks to all participating children and their families for their contribution to this study.

## References

Clinical and Laboratory Standards Institute (CLSI). Blood Gas and pH Analysis and Related Measurements; Approved Guideline—Second Edition. CLSI document C46-A2. Wayne, PA: Clinical and Laboratory Standards Institute; 2009.

Diniz, E.T., et al., 2013. Primary hyperparathyroidism is associated with subclinical peripheral neural alterations. Endocr. Pract. 19, 219–225.

Farrar, M.A., et al., 2013. Evolution of peripheral nerve function in humans: novel insights from motor nerve excitability. J. Physiol. 591, 273–286.

Fleming, L.W., Lenman, J.A., Stewart, W.K., 1972. Effect of magnesium on nerve conduction velocity during regular dialysis treatment. J. Neurol. Neurosurg. Psychiatry 35, 342–355.

Frankenhaeuser B. The effect of calcium on the myelinated nerve fibre. J Physiol. 1957;137(2):245–260. doi:10.1113/jphysiol.1957.sp005809. PMID: 13449875.

Freeman, S.A., Desmazières, A., Fricker, D., Lubetzki, C., Sol-Foulon, N., 2016. Mechanisms of sodium channel clustering and its influence on axonal impulse conduction. Cell. Mol. Life Sci. 73, 723–735.

García, A., Calleja, J., Antolín, F.M., Berciano, J., 2000. Peripheral motor and sensory nerve conduction studies in normal infants and children. Clin. Neurophysiol. 111, 513– 520.

Gerhart, D.Z., Drewes, L.R., 1990. Glucose transporters at the blood–nerve barrier are associated with perineurial cells and endoneurial microvessels. Brain Res. 508, 46–50.

Goldstein, D.A., Chui, L.A., Massry, S.G., 1978. Effect of parathyroid hormone and uremia on peripheral nerve calcium and motor nerve conduction velocity. J. Clin. Invest. 62, 88–93.

Harrington, M.G., et al., 2006. Cerebrospinal fluid sodium increases in migraine. Headache 46, 1128–1135.

Hille B. Charges and potentials at the nerve surface. Divalent ions and pH. J Gen Physiol. 1968;51(2):221–236. doi:10.1085/jgp.51.2.221.

Holm S. A simple sequentially rejective multiple test procedure. Scand J Stat. 1979;6(2):65–70

Husted, R.F., Reed, D.J., 1976. Regulation of cerebrospinal fluid potassium by the cat choroid plexus. J. Physiol. 259, 213–221.

Jenny, C., Lütschg, J., Broser, P.J., 2020. Change in cross-sectional area of the median nerve with age in neonates, infants and children analyzed by high-resolution ultrasound imaging. Eur. J. Paediatr. Neurol. 29, 137–143.

Kiernan, M.C., Walters, R.J.L., Andersen, K.V., Taube, D., Murray, N.M.F., Bostock, H., 2002. Nerve excitability changes in chronic renal failure indicate membrane depolarization due to hyperkalaemia. Brain 125, 1366–1378.

Maurer, K., Bostock, H., Koltzenburg, M., 2012. Protons regulate the excitability properties of rat myelinated sensory axons in vitro through block of persistent sodium currents. J. Peripher. Nerv. Syst. 17, 102–111.

Mert, T., Gunes, Y., Guven, M., Gunay, I., Ozcengiz, D., 2003. Effects of calcium and magnesium on peripheral nerve conduction. Pol. J. Pharmacol. 55, 25–30.

Myers, R.R., Heckman, H.M., Powell, H.C., 1983. Endoneurial fluid is hypertonic. Results of microanalysis and its significance in neuropathy. J. Neuropathol. Exp. Neurol. 42, 217–224.

Oberhauser, S.S., l’Allemand, D., Willems, E.P., Gozzi, T., Heldt, K., Eilers, M., Stasinaki, A., Lütschg, J., Broser, P.J., 2023. Slowing of peripheral nerve conduction velocity in children and adolescents with type 1 diabetes is predicted by glucose fluctuations. Diabetes 72, 1835–1840.

Ørskov, L., Worm, M., Schmitz, O., Mengel, A., Sidenius, P., 1994. Nerve conduction velocity in man: influence of glucose, somatostatin and electrolytes. Diabetologia 37, 1216–1220.

Palladino, S.P., et al., 2017. The human blood–nerve barrier transcriptome. Sci. Rep. 7, 17477.

Parano, E., Uncini, A., De Vivo, D.C., Lovelace, R.E., 1993. Electrophysiologic correlates of peripheral nervous system maturation in infancy and childhood. J. Child Neurol. 8, 336–338.

Poliak, S., Peles, E., 2003. The local differentiation of myelinated axons at nodes of Ranvier. Nat. Rev. Neurosci. 4, 968–980.

Rasband, M.N., Peles, E., 2016. The nodes of Ranvier: molecular assembly and maintenance. Cold Spring Harb. Perspect. Biol. 8, a020495.

Rechthand, E., Smith, Q.R., Rapoport, S.I., 1989. Structural specificity of sugar transport at the blood–nerve barrier. J. Neurochem. 53, 119–123.

Saunders, N.R., et al., 2016. Fluid and ion transfer across the blood–brain and blood– cerebrospinal fluid barriers: a comparative account of mechanisms and roles. Fluids Barriers CNS 13, 19.

Schwarz, J.R., 2021. Function of K2P channels in the mammalian node of Ranvier. J. Physiol. 599, 4895–4908.

Ubogu, E.E., 2013. The molecular and biophysical characterization of the human blood– nerve barrier: current concepts. J. Vasc. Res. 50, 289–303.

Weerasuriya, A., 1987. Permeability of endoneurial capillaries to K, Na and Cl and its relation to peripheral nerve excitability. Brain Res. 419, 188–196.

Z’Graggen, W.J., Lin, C.S.Y., Howard, R.S., Beale, R.J., Bostock, H., 2006. Nerve excitability changes in critical illness polyneuropathy. Brain 129, 2461–2470.

Zhang, Q., et al., 2018. Low serum phosphate and magnesium levels are associated with peripheral neuropathy in patients with type 2 diabetes mellitus. Diabetes Res. Clin. Pract. 146, 1–7.

Zhang, Z., David, G., 2016. Stimulation-induced Ca²⁺ influx at nodes of Ranvier in mouse peripheral motor axons. J. Physiol. 594, 39–57.

